# Hypothesizing mechanistic links between microbes and disease using knowledge graphs

**DOI:** 10.1101/2023.12.01.569645

**Authors:** Brook Santangelo, Michael Bada, Lawrence Hunter, Catherine Lozupone

**Affiliations:** Computational Bioscience Program, University of Colorado Denver Anschutz Medical Campus Aurora, CO 80045, USA

**Keywords:** knowledge graphs, embedding methods, microbiome

## Abstract

Knowledge graphs have found broad biomedical applications, providing useful representations of complex knowledge. Although plentiful evidence exists linking the gut microbiome to disease, mechanistic understanding of those relationships remains generally elusive. Here we demonstrate the potential of knowledge graphs to hypothesize plausible mechanistic accounts of host-microbe interactions in disease. To do so, we constructed a knowledge graph of linked microbes, genes and metabolites called MGMLink. Using a semantically constrained shortest path search through the graph and a novel path prioritization methodology based on cosine similarity, we show that this knowledge supports inference of mechanistic hypotheses that explain observed relationships between microbes and disease phenotypes. We discuss specific applications of this methodology in inflammatory bowel disease and Parkinson’s disease. This approach enables mechanistic hypotheses surrounding the complex interactions between gut microbes and disease to be generated in a scalable and comprehensive manner.

## 1. Introduction

That gut microbiome composition differs in disease states has become increasingly clear from metagenomic studies [1,2]. However, our understanding of specific microbiome signatures on clinical outcomes is sparse and mainly correlative [3,4]. Microbiome signatures have been associated with auto-immune, gastrointestinal, cancer, and neurological disease [5,6], but the conclusions often lack a mechanistic account. Experimental studies to test hypothetical mechanisms are critical, but they are expensive and cannot cover all potential pathways a microbe might act on within a host. Additionally, investigations of immune and metabolomic outcomes in relation to microbial abundance lack a comprehensive integration of biomedical knowledge. Without this mechanistic account we do not fully understand the role of the gut microbiome in disease, which makes the development of better targeted therapeutics more difficult.

With more than 20,000 citations relating the microbiome to disease now published each year in PubMed since 2019, there is huge potential for integrating existing studies into a centralized knowledge base [7]. The complexity and multi-omic nature of the gut microbiome requires a comprehensive view of how microbes and their metabolic products interact with the human body. We therefore seek to provide contextual insight to predict how these interactions influence disease. To address the limitations of correlative analysis, we present a methodology that integrates microbiome information into knowledge graphs (KGs) and generates hypothetical mechanistic accounts of how microbes influence disease. KGs describe relationships between entities of different types, *e*.*g*., how a gene or gene product influences a disease, or how a metabolite is processed in a specific pathway. KGs have broad applications across biomedical research, including prediction of drug-drug interactions [8,9], evaluation of mechanisms of toxicity [10], prediction of unknown drug disease targets [11], or linking symptoms from electronic health records to better understand disease inference [12]. However, these applications have yet to be extended into the microbiome field [7]. Existing microbial KGs incorporate information more focused on microbial trait outcomes [13] or are limited in scope and/or lacking in biomedical information [2,14,15]. Here we describe the creation and application of a KG of microbe-host interactions toward the creation of mechanistic hypotheses regarding microbe-disease relationships.

We created a microbiome relevant KG representing microbe-gene-metabolite links called MGMLink by integrating known microbe-host interactions into a biomedical KG built using the PheKnowLator system (Figure 1). PheKnowLator is a Python 3 library that enables construction of KGs that incorporate a wide variety of data and terminology sources, including ontologies such as the Monarch Disease Ontology (MONDO), the Chemicals Entities of Biological Interest Ontology (CHEBI), and the Human Phenotype Ontology (HPO) [16]. To augment the default PheKnowLator KG with information on microbes, we integrated data from gutMGene, a manually curated repository of assertions involving gut microbes, microbial metabolites, and target genes from over 360 PubMed publications [17]. Using MGMLink, we introduce a method for examining mechanisms that describe the interaction between a microbe and a target entity. We evaluate the utility of this KG in the discovery of microbial mechanisms by exploring the extent to which microbes included in the KG are relevant in specific diseases (inflammatory bowel disease and Parkinson’s disease) and demonstrating two feasible mechanisms of action between microbes and these diseases. We thus show that the combination of knowledge from gutMGene and PheKnowLator facilitates hypothesis generation regarding mechanisms linking microbes with disease.

**Figure 1.**
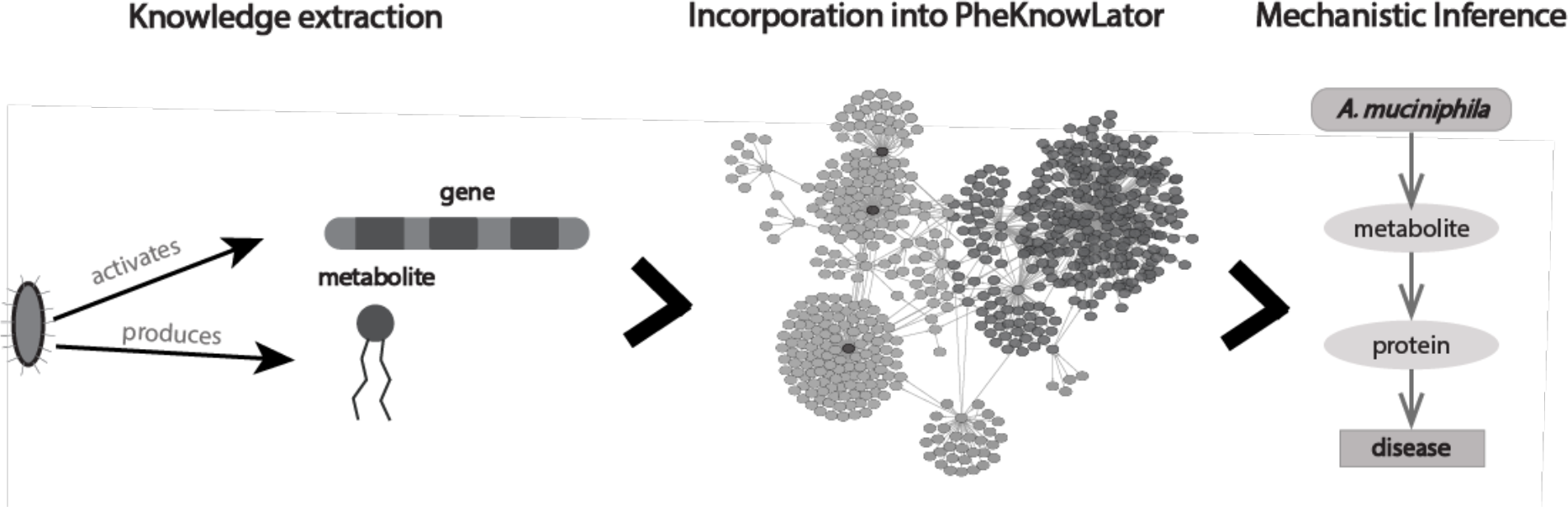
Microbe-host gene and microbe-metabolite relationships such as the types shown (microbe activates gene or microbe produces metabolite) using the gutMGene database were incorporated into the PheKnowLator framework. The microbiome-relevant KG was then used to examine paths between microbes and diseases of interest.

## 2. Methods

### 2.1 Knowledge Representation and Incorporation

To generate MGMLink, we incorporated previously published microbe-host interactions from the gutMGene database into the PheKnowLator framework. The framework allows alternative knowledge modeling approaches; we used a model in which concepts are unidirectionally relationally linked in the graph (as opposed to bidirectionally) [16]. We also use a version of a PheKnowLator KG that reflects a transformation based on OWL-NETS, which enables network inference of OWL-encoded knowledge via abstraction into biologically meaningful triples [18]. These parameters produce the topologically simplest graph and has the best performance in relevant metrics such as node embedding quality. The representation of gutMGene data in the framework was matched to the PheKnowLator framework parameters. All assertions were mapped to an OWL-encoded KG representation and then transformed into triples using OWL-NETS, resulting in four unique patterns, examples of which are shown in Table 1 [18]. This resulted in new KG nodes that represented microbes in the context of the anatomical location (*i*.*e*., the gut) and species (human or mouse) in which the interactions have been reported to occur.

**Table 1.**
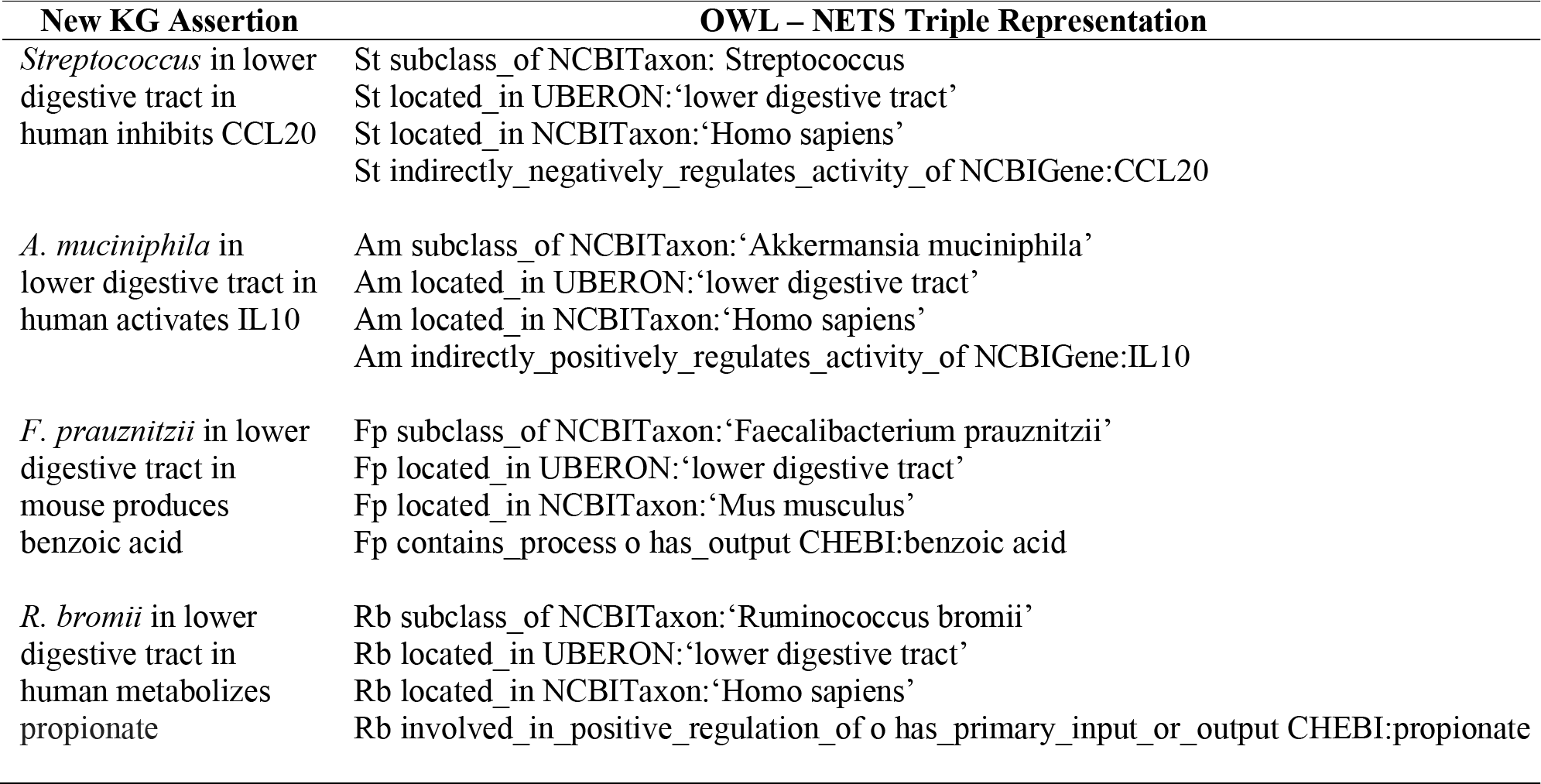
Specific examples of all types of patterns identified from gutMGene assertions, and corresponding OWL-NETS representations into which these assertions were converted to integrate them into the knowledge base.

The gutMGene database consists of microbe-metabolite and microbe-gene assertions that occur in the host of either human or mouse. These relationships were manually extracted from over 360 PubMed publications and are based on validated methods such as RT-qPCR, high-performance liquid chromatography, and 16S rRNA sequencing [17]. Specifically, the gutMGene database describes four different types of assertions: (1) microbial substrates observed in humans or mice, (2) microbial metabolites observed in humans or mice, (3) genes observed to be inhibited by microbes in humans and mice, and (4) genes observed to be activated by microbes in humans and mice [17]. Assertions 1 and 2 were extracted from the Association between Gut microbe and Metabolite v1.0 results for Human and Mouse, respectively, and assertions 3 and 4 were extracted from the Association between Gut microbe and Gene v1.0 for Human and Mouse, respectively. These were then represented using a specific semantic pattern to encompass the relationship type and the context (Table 1). We attempted to map each microbial type to an entry in the NCBI Taxonomy, and for those microbial types that could not be mapped to NCBI Taxonomy entries, new nodes representing unclassified bacterial organisms were created. Additionally, microbes that had an unclear or indirect mapping to a NCBITaxon identifier were corrected. A total of 1874 assertions from gutMGene were created in MGMLink for 533 unique microbial taxa (at the family, genus, species, or strain level). NCBI Taxonomy entries were found for 336 of these taxa, and contextual information is incorporated for 461 of the gutMGene assertions. To generate a connected graph, species or strains were related to their corresponding higher-level classifications (genus, family, class, etc.) which already existed in the KG. This allowed for inferences to be made regarding species-strain or genus-species interactions.

### 2.2 Mechanism Prediction Framework

The path between two nodes in a KG can be drawn in a nearly infinite number of ways, especially for a KG of this size (over 780,000 nodes and 5,000,000 edges). Here we employ a shortest-path search using a breadth-first search algorithm with unweighted edges, which incrementally searches all neighbors of a source node until the target node is found and returns the path with the minimum number of edges. Shortest path search is a common approach to examining a mechanistic link between two nodes [19,20]. The result is unbiased, allowing for a comprehensive assessment of potential interactions between microbes and metabolites, proteins, or processes of interest. Exclusion of specific semantic constraints, e.g., “only_in_taxon Homo sapiens”, or “part_of Homo sapiens”, remove paths likely to be irrelevant to the mechanistic accounts we are seeking.

The source inputted into this process can represent a microbe or a first-order neighbor of a microbe. Using the first-order neighbors of microbes as the source node in the search allows for more potential mechanistic explanations of a microbial interaction to be hypothesized. To semantically constrain this search, we only included host genes or microbial metabolites as the first-order neighbors as the source of the shortest path search (Figure 2).

**Figure 2.**
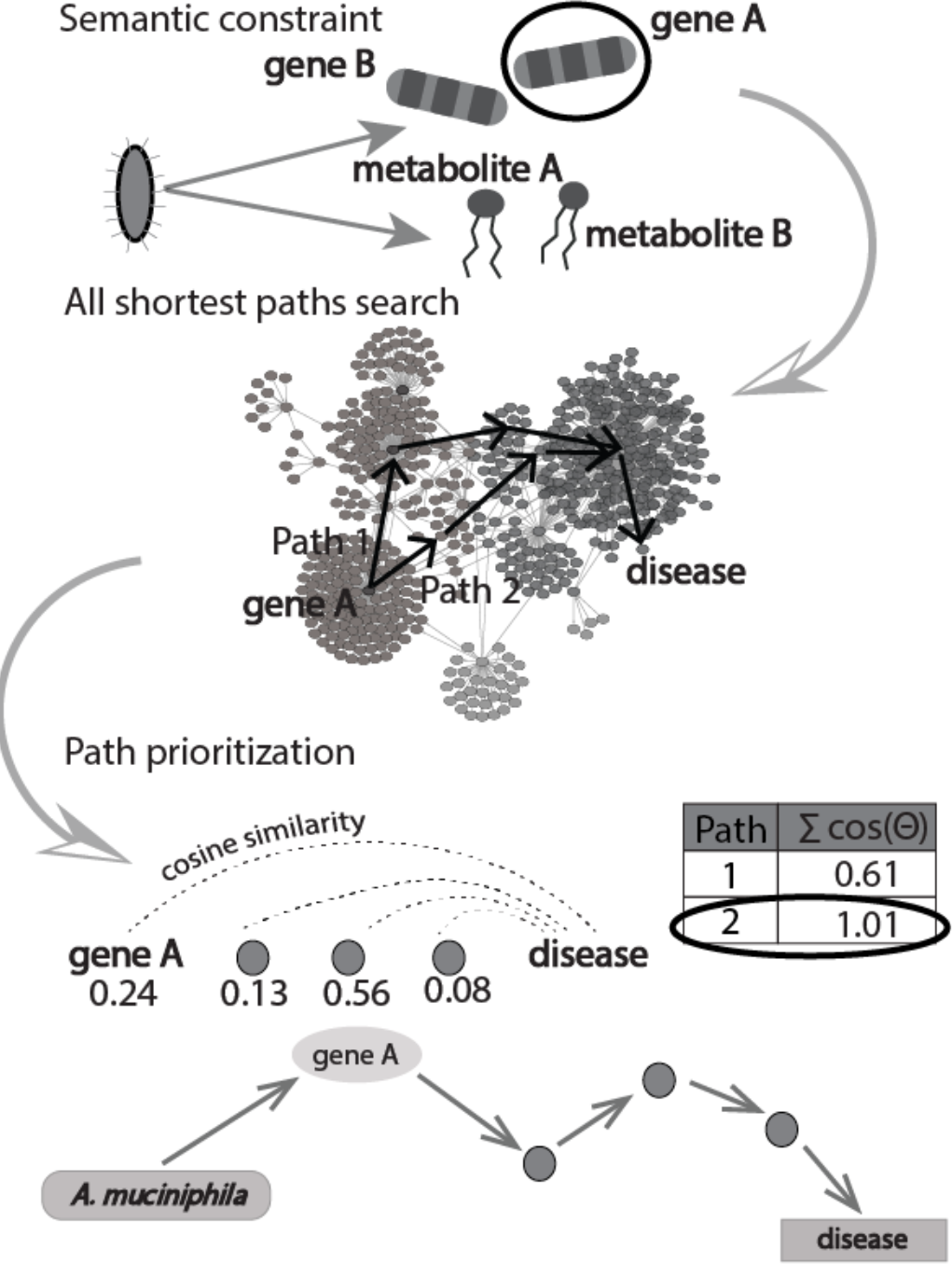
Methodology used to prioritize paths between a given microbe and disease of interest.

There are frequently many ties among shortest paths, so we devised a novel metric to prioritize among these paths. For each path, the total cosine similarity between all nodes (source node and all intermediate nodes) in the path is compared to the target node and the path with the closest value was chosen (Figure 2). Vector embeddings of MGMLink were generated using Node2Vec [21]. The dimensions of the vector embeddings were selected by minimizing the information loss while also minimizing the dimensions. We applied the methodology used by Grover et al., which evaluates the information loss by calculating the macro F1-score for the multi-label classification of nodes in the graph according to their ontology or database (out of 51 possible labels) and identifying the point of inflection where no significant improvement in labeling occurs with more dimensions [21]. By maximizing total cosine similarity among all shortest paths, the path search is biased towards nodes that have the most similar neighbors, which may prioritize more relevant paths (Figure 2).

## 3. Results

To assess the quality of MGMLink and the prioritized shortest path approach to hypothesizing mechanisms, we explored the two representative diseases with known microbial mechanisms, inflammatory bowel disease or Parkinson’s disease. Many studies have identified changes in the microbiome for individuals with inflammatory bowel disease (IBD) as compared to healthy individuals, citing significantly decreased abundances of Prevotella, Faecalibacterium, Clostridium, Bifidobacterium and increased abundances of Fusobacterium and Gardnerella [22–24]. However, we are in the early stages of exploring the mechanisms of microbial influence in IBD [25–28]. Parkinson’s disease (PD) is the second most prevalent neurological disease and is associated with motor, gastrointestinal, and psychiatric dysfunction [29]. It is unknown whether PD starts in the gut, though integrated understanding of intestinal permeability and the pathological implications of the disease highlight the importance of the gut microbiome in the development of PD. Bacteria can produce neurotransmitters and neuromodulators that may affect the pathogenesis of the disease, and the differences in microbiome signatures have become clearer in healthy vs PD individuals [14]. As shown in Figure 3, there are an abundance of paths between these microbes with those diseases in MGMLink. This finding reiterates the importance of our methodology to prioritize all shortest paths between two nodes of interest.

**Figure 3.**
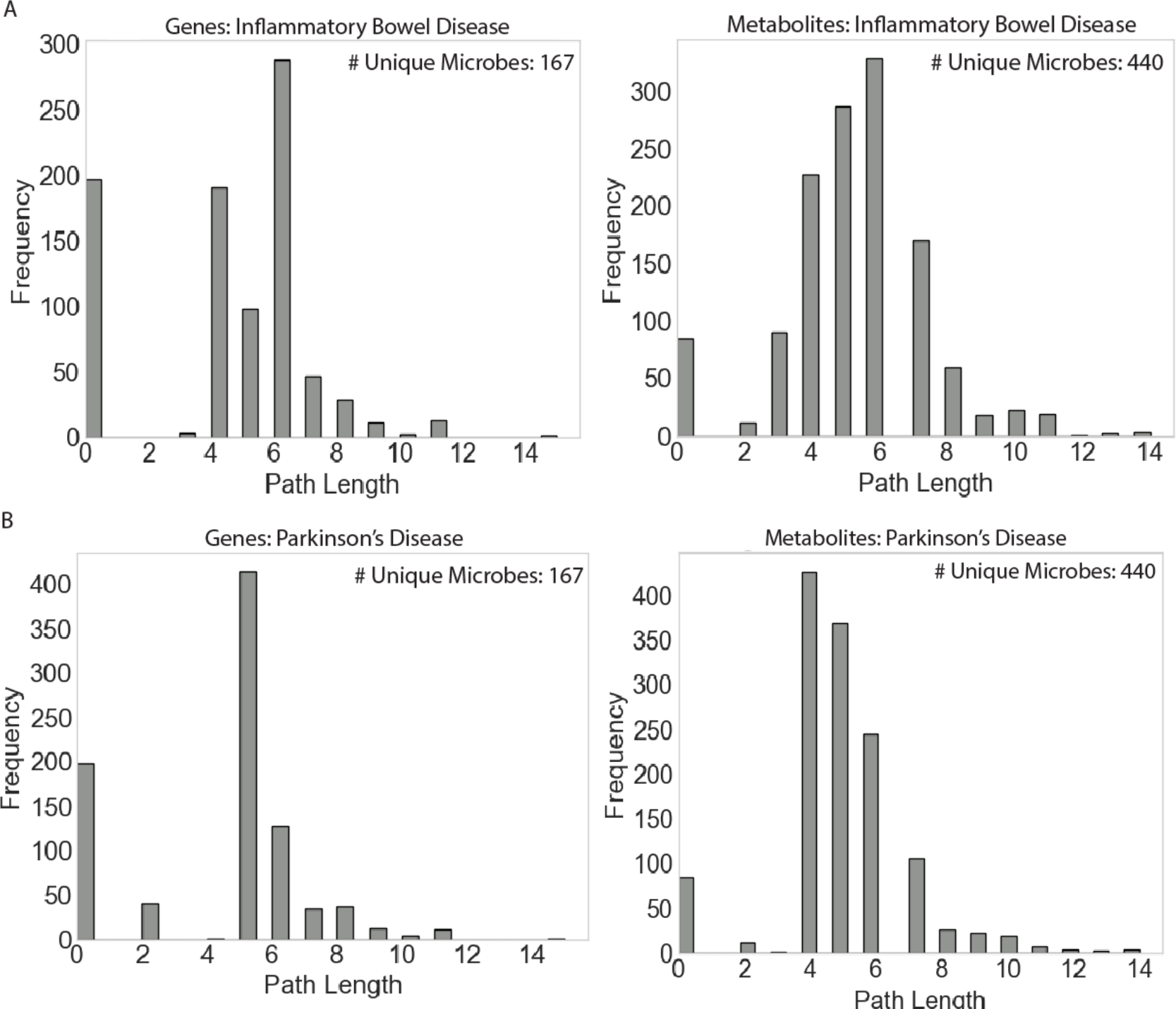
All genes or metabolites that were found to have a potential path through the graph to IBD (A) or PD (B). The left panel shows the frequency of path lengths that went through a microbe-host gene interaction, and the right through a microbe-metabolite interaction. A path length of 0 represents a case where no path could be found between an entity and the corresponding disease. The number of microbes from which a path could be drawn to the corresponding disease through a gene or metabolite is also reported.

### 3.1. Inflammatory Bowel Disease

We next aimed to determine whether we could expand upon previous observations using MGMLink. We applied this knowledge-based methodology to a study that compared individuals with and without IBD using host gene expression and 16S microbial relative abundances [30]. This allowed us to determine whether our knowledge base could provide more detailed paths between microbe: gene: IBD relationships than would be evident from this study alone. IBD is characterized by chronic inflammation of the gut, where excess cytokine production in the mucosa results in gastrointestinal discomfort [31]. Generally, the pathogenesis of IBD is thought to be driven by dysbiosis of the microbiome that influences an abnormal immune response, though the direct cause of key microbes in the disease is unknown [25]. We examined the shortest path between microbes of interest and IBD to hypothesize potential mechanisms underlying differentially abundant microbes in IBD and the pathogenesis of the disease.

The multi-omic study, conducted as part of the Integrative Human Microbiome Project, obtained gene expression and microbiome signatures of 132 individuals with or without IBD over the course of one year through sampling of stool, biopsy, and blood [30]. Significantly differentially expressed genes identified from biopsies in the ileum and rectum were compared to differentially abundant microbes identified through 16S sequencing of identical biopsy samples based on partial Spearman correlation that accounted for BMI, age, sex, and diagnosis. In total, 178 gene-microbe pairs that co-varied and were significantly different among IBD and non-IBD individuals were identified [30]. The gutMGene database only contained 12 out of 40 represented microbes from the gene-microbe pairs, alluding to the incomplete nature of MGMLink which limits its application. 3 of the 178 gene-microbe pairs existed in the knowledge base allowing for further interpretation of the suggested microbe-host interaction (Table 2). These microbe-gene relationships that exist in gutMGene were identified in the same IBD analysis over which this evaluation was done. Although confirmatory evidence of this interaction is lacking in MGMLink, its presence suggests that this knowledge base can make cited studies more accessible. More interestingly this methodology uncovered potential mechanistic explanations for this gene-microbe-disease interaction that were previously unknown.

**Table 2.** Gene-microbe pairs from gene expression and microbial abundance analysis that exist in MGMLink.

| Microbe | Gene | Regulation |
| --- | --- | --- |
| Eubacterium rectale (human) | CXCL6 | Negative |
| Streptococcus (human) | CCL20 | Negative |
| Eikenella (human) | CCL20 | Negative |

We identified all first order neighbors of each of the microbes of interest (*Eubacterium rectale, Streptococcus*, and *Eikenella*), and evaluated the shortest path between that neighbor and IBD (Figure 2b). *Streptococcus* was found to be negatively correlated (R2 = -0.53) with CCL20 by this analysis [30], and that microbe-gene relationship was represented semantically as shown in Figure 2a. The result of evaluating the shortest path between CCL20 and IBD was a hypothesized mechanism that involved C-C motif chemokine 20, IL-1β, and prostaglandin E synthase (Figure 4). A previous study cited CCL20 as an inducer of IL-1β, a pro-inflammatory cytokine that is increased in IBD and results in the chronic inflammation that is a staple of the IBD phenotype [25]. This was done by evaluating cytokine signatures of peripheral blood mononuclear cells extracted from individuals with IBD, suggesting a clear association between CCL20 and IL-1β [25]. Additionally, another study which used a mouse model of Ulcerative Colitis (UC), a class of IBD, to evaluate the role of prostaglandins, a known inhibitor of the inflammatory response, in the phenotype observed from UC found that prostaglandin E2 (PGE2) plays a significant role in intestinal homeostasis [32]. PGE2 synthesis is mediated by multiple enzymes, including microsomal prostaglandin E synthase which is included in this path identified. *Streptococcus pneumoniae* was cited as a pathogenic microbe that can stimulate IL-1β secretion which is further boosted by PGE2 signaling [33]. PGE2 signaling can be through 4 different G-protein coupled receptors resulting in differing immune signaling degradation into an inactive form thus the exact mechanism at play here is unknown [33], however these studies further support the plausible mechanism predicted in Figure 4. Although these results have yet to be validated with an experimental approach, the potential for generating targets in a high throughput manner is clear and this method reduces the search space of microbial targets.

**Figure 4.**
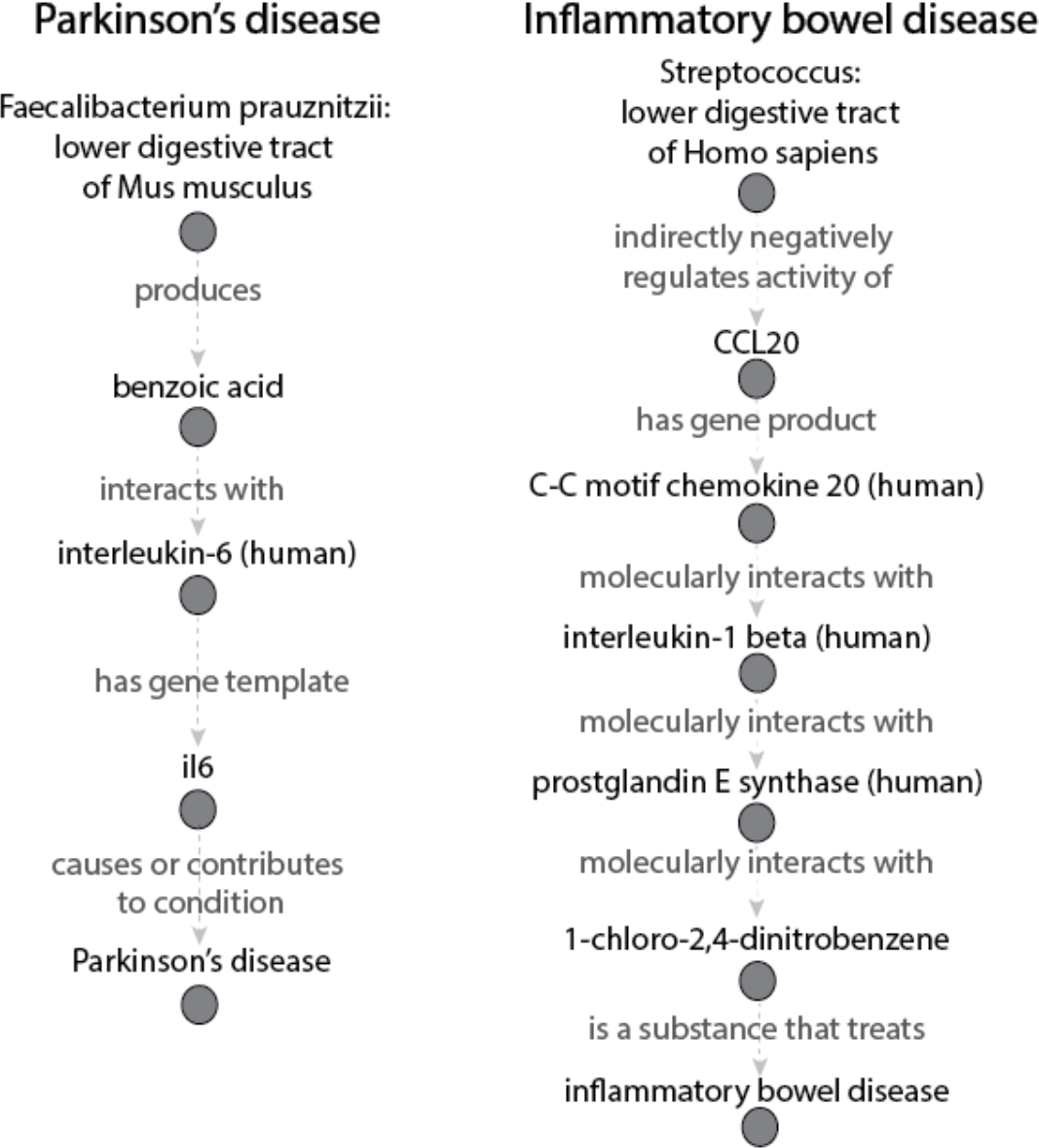
Shortest path results for searching all first order neighbors of *Faecalibacterium prausnitzii* as the source and PD as the target (left) and *Streptococcus* as the source and IBD as the target (right).

### 3.2. Parkinson’s Disease

The next case study identified a microbe associated with PD that was previously unknown in the context of the experiment, *Faecalibacterium prauznitzii. F. prauznitzii* is known to have a protective effect for PD [29]. The cosine similarity was calculated between PD and each microbe that exists in the graph to identify *F. prauznitzii* as the most closely associated microbe to the disease. This relationship was then expanded on with a potential mechanism of the microbe’s protective effects. Although there are limitations in this methodology with a biased and incomplete knowledge base, it is promising that we were able to identify a prominent microbe to be associated with PD.

An all-shortest paths search was then conducted between *F. prausnitzii* and PD, again evaluating the path through each first order neighbor of *F. prausnitzii* as the first step in the path. We prioritized the paths for each first order node-disease pair using the total cosine similarity (Figure 2b). This resulted is a hypothesized mechanism suggesting that a metabolite produced by *F. prausnitzii* interacts with the pro-inflammatory cytokine IL-6 (Figure 4). Benzoic acid has been seen to have anti-inflammatory effects [34–36], and *F. prausnitzii* has been observed to be decreased in patients with PD [37]. It is plausible that a reduction in the anti-inflammatory microbe *F. prauznitzii* may be implicated in the inflammatory state of a person with PD through the lack of inhibition of the pro-inflammatory cytokine IL-6.

## 4. Discussion and Conclusion

Knowledge graphs have extensive applications in the biomedical field because they integrate complex concepts in a systematic way. Network representations of KGs have enabled domain experts to better understand potential interactions between genes, phenotypes, drugs, and diseases in a high throughput manner, allowing for expensive wet lab experiments to be more focused and informed. KGs have been built for more comprehensive gene prioritization which can enhance knowledge of drug-gene interactions or describe gene targets for a specific disease [38,39]. KGs have also been used to understand drug-drug interactions or hypothesize drug repurposing directions [10,40]. Furthermore, there is potential to expand the scientific reach of these methodologies into the field of the microbiome. To the best of our knowledge, the microbiome relevant MGMLink KG described here is the first to realize this potential.

The computational power of networks analysis techniques can integrate information from thousands of publications to allow the use of network algorithms to generate hypotheses about microbes influence diseases such as inflammatory bowel disease and Parkinson’s disease. This method identified an implicit microbe-disease relationship in PD that was validated with existing literature. Additionally, this methodology expanded upon previously observed microbe-gene interactions with a mechanistic explanation. These results demonstrate that a semantically constrained shortest path search presents feasible descriptions of the ways a microbe may influence disease.

This work introduces a method for using novel representations of existing knowledge about microbial function to hypothesize mechanisms. However, there limitations to this work. MGMLink contains only the limited number of microbes described in the gutMGene database and does not cover all mechanisms by which microbes influence host genes or consume or produce metabolites. Additionally, the shortest path search methodology is only one example of networks analysis techniques that can be used to examine the graph. We are focused on more extensive semantic restrictions of the network space that can be informed by domain experts and lead to useful mechanistic hypotheses. Lastly, using literature review as an evaluation of these results lacks robustness and limits us from predicting unknown, novel hypotheses. In the future, we foresee these methods proposing hypothesized microbe-host interactions that bring researchers closer to targeted lab experimentation. Despite these limitations, this work offers a novel approach to addressing the pressing need to understand the mechanisms underlying the now well-established relationships between gut microbes and human disease.

